# Organ-confined mRNA delivery and protein expression during *ex vivo* perfusion of porcine donor kidneys

**DOI:** 10.1101/2025.08.28.672793

**Authors:** Cristina Ballester, Ping Song, Katerina B Knudsen, Chris Jaynes, Henri Leuvenink, Victor R Llorente, Annick Van Furth, Henrik Hager, Rikke Nørregaard, Jan-Luuk Hillebrands, Anna Krarup Keller, Bente Jespersen, Jørgen Kjems, Marco Eijken

## Abstract

Achieving efficient mRNA delivery and expression in intact solid organs beyond the liver remains a major challenge for advancing mRNA therapeutics. Normothermic machine perfusion (NMP) provides a clinically relevant *ex vivo* platform for targeted intervention during donor organ preservation, minimizing systemic exposure and enabling organ-confined mRNA delivery. Here, we demonstrate that intra-arterial infusion of lipid nanoparticles (LNPs) encapsulating reporter mRNAs during kidney NMP enables robust, organ-wide protein expression in human-sized porcine donor kidneys. Engineered LNP formulations were first evaluated for stability and efficacy in kidney cell lines, where LNPs pegylated with TPGS (D-α-tocopherol polyethylene glycol 1000 succinate) achieved the highest reporter protein expression levels. Next, the LNP-encapsulated mRNAs were introduced into porcine kidneys via a 5-minute infusion during NMP. mCherry and human erythropoietin (hEPO) mRNAs were delivered via the renal artery, serving as intracellular and secreted reporters, respectively. The perfusion was continued for 6-12h after LNP-mRNA infusion with the erythrocyte-based perfusate at 37 °C. LNP-mRNA administration resulted in widespread parenchymal mRNA uptake and rapid, robust reporter protein expression that continued to increase over the perfusion period. hEPO protein was detected in perfusate and urine, while mCherry expression localized to endothelial and tubular cells within the renal parenchyma. Consistent with the *in vitro* data, TPGS-LNPs produced the highest expression during NMP. Importantly, LNP-mediated mRNA delivery during NMP did not affect perfusion parameters or histological integrity. These findings demonstrate successful mRNA-driven protein expression in intact donor kidneys, establishing NMP as a clinically relevant route for organ-specific RNA delivery and protein expression modulation.

**One sentence summary:** Using a clinically compatible normothermic machine perfusion model, we demonstrate that lipid nanoparticle-mediated mRNA delivery enables rapid, organ-wide protein expression in human-sized porcine donor kidneys.

## INTRODUCTION

Recent advancements in mRNA-based therapeutics have significantly expanded their potential applications in healthcare. The clinical success of SARS-CoV-2 mRNA vaccines and recent progress in mRNA cancer vaccines highlight the versatility of this platform(1–3). Beyond vaccines, synthetic mRNAs are being developed for therapeutic protein replacement, immunotherapies, genome editing and tissue repair(4). Numerous clinical trials exploring mRNA therapeutics are ongoing across a wide range of diseases, including cystic fibrosis transmembrane conductance regulator (CFTR) mRNA for cystic fibrosis and vascular endothelial growth factor A (VEGF-A) mRNA for myocardial infarction(5, 6), reflecting the growing therapeutic potential of this platform. Delivery of therapeutic mRNAs requires safe, effective and stable delivery systems to avoid nucleic acid degradation and overcome biological barriers. Lipid nanoparticles (LNPs) have been widely used as an efficient delivery platform to enhance mRNA stability and cellular uptake(7, 8). For non-vaccine applications, therapeutic mRNAs typically require higher dosing, alternative routes of administration beyond intramuscular injections, and formulation strategies for enhanced targeted delivery and functional expression. In the bloodstream, certain LNPs can bind to apolipoprotein E (APOE), which, in turn, interacts with the abundant low-density lipoprotein receptors (LDLRs) in the liver. This interaction promotes preferential uptake by hepatocytes and limits distribution to other tissues(9, 10) Accordingly, local delivery routes such as direct injection or nebulization have been explored to improve organ-specific delivery, yet achieving consistant and widespread distribution within a complex organ remains challenging(5, 11). An alternative strategy is e*x vivo* organ machine perfusion, a technique that circulates an oxygenated preservation solution or immune cell-depleted blood through the organ’s vascular network to maintain its function and viability, which is increasingly adopted in clinical organ transplantation practice(12–14). The perfusion can be performed under hypothermic conditions (typically 4-10°C), which minimizes metabolic activity and cellular damage, or under normothermic conditions (35–37°C), closely mimicking physiological conditions and metabolic rates. This dynamic organ preservation technique offers a unique opportunity for *ex vivo* delivery of therapeutics to repair or improve the organ prior to transplantation. Despite advances in transplantation, the persistent shortage of suitable donor organs and the frequent presence of ischemic or other injury-related damage continue to limit graft function and long-term survival, underscoring the need for novel strategies to improve donor organ quality before transplantation(15–17). Delivering synthetic mRNAs during machine perfusion enables controlled, organ-wide delivery with minimal off-target exposure, which remains a critical barrier in the mRNA field. In particular, normothermic machine perfusion (NMP) supports active metabolism, which can facilitate LNP uptake and initiate mRNA translation prior to transplantation of the organ. PEG-lipids strongly influence LNP biophysical and biological behavior, including stability in biological fluids and interactions with tissues(18, 19). Our previous work demonstrated that altering the PEG-lipid structure modulates mRNA delivery efficiency, making PEG-lipid chemistry a rational parameter to optimize for ex vivo mRNA delivery(20). In this study, we evaluated the feasibility of the *ex vivo* delivery of therapeutic mRNAs during NMP. Reporter mRNAs were included to assess delivery efficiency, spatial distribution, and biosafety, providing a foundation for mRNA-based therapies to restore and enhance donor organ function.

## RESULTS

### Preparation and characterization of LNPs

In the present study, we developed three LNP formulations based on our in-house synthesized ionizable lipid, C14120 (Supplemental figure 1), using different coating materials: PEG-lipid TPGS (TPGS-LNP), PEG-lipid DSPE-PEG2000 (DSPE-LNP), and a PEG-based block copolymer (F127-LNP). The LNPs ranged approximately between 150 nm to 175 nm in diameter and exhibited a positive surface charge with zeta potential over 30 mV at pH 5.5 and generally low polydispersity (<0.2) (Figure 1a-c). LNPs were loaded with mRNA through electrostatic interactions between ionizable lipids and negatively charged mRNAs. mRNAs were loaded freshed right before the treatment both *in vitro* and *ex vivo*. mRNA encoding mCherry protein was encapsulated within the three LNP formulations at N/P ratios (the stoichiometry between protonatable nitrogen atoms (N) in ionizable lipid and anionic phosphor atoms(P) in a nucleic acid) of 6, 12, and 16 and evaluated for translational output in OK cells.

**Figure 1:**
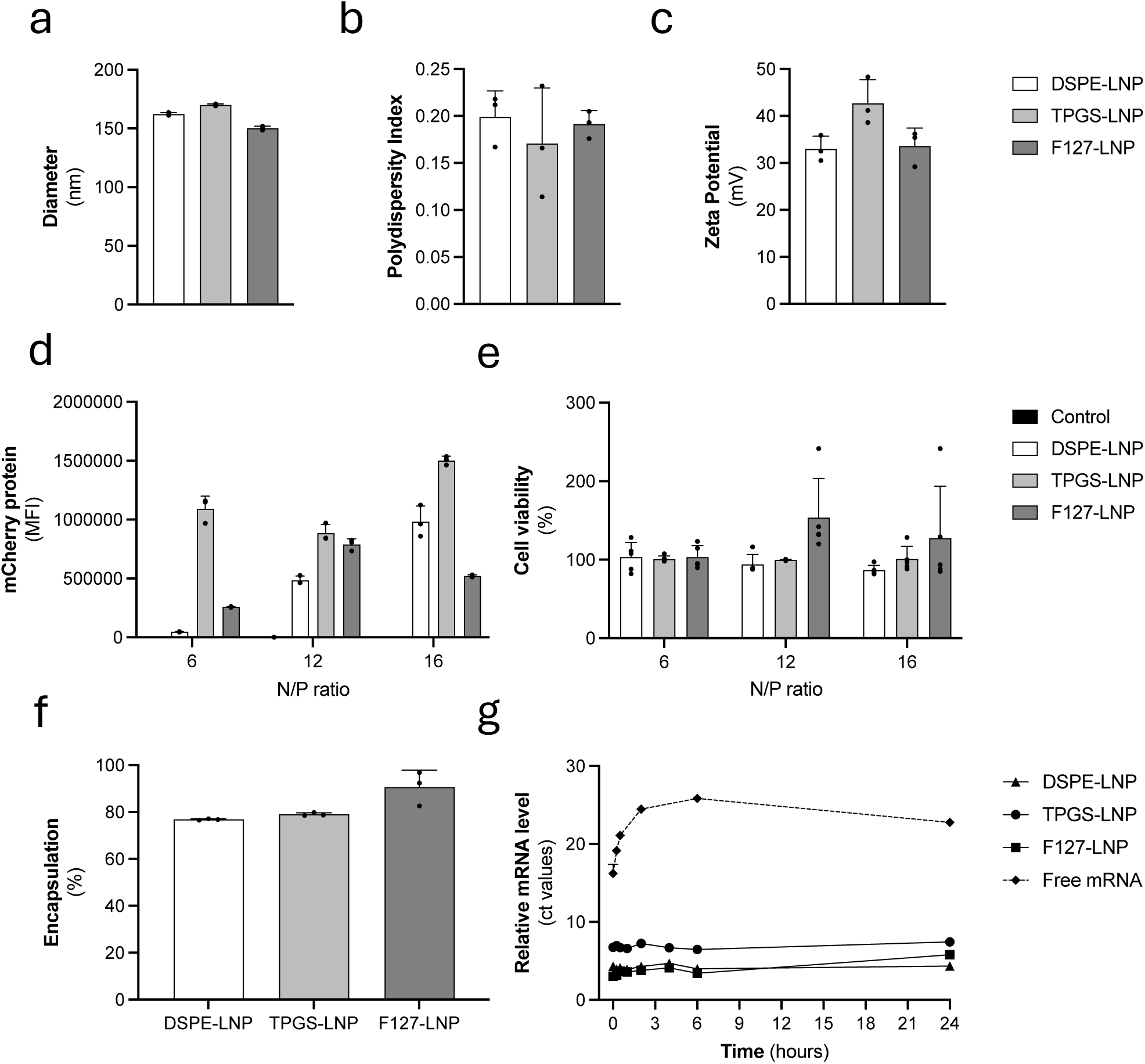
In vitro Characterization of F127-LNP, TPGS-LNP and DSPE-LNP. (a) Hydrodynamic diameter. (b) Polydispersity index (PDI). (c) Zeta potential. (d) mCherry protein expression and cell viability (e) in OK cells after transfection with LNP-encapsulated mCherry mRNA at N/P ratios of 6, 12 and 16. (f) The encapsulation efficiency of F127-LNP, TPGS-LNP and DSPE-LNP at N/P ratio of 12. (g) Stability of free and LNP-encapsulated mCherry mRNA (F127-LNP, TPGS-LNP and DSPE-LNP) in kidney perfusion solution over 24 hours at 37 °C, indicated by ct values measured by RT-qPCR and presented as Mean ± SD (n = 3).

DSPE-LNP showed an increase in expression with rising N/P ratios whereas the F127-LNP formulation peaked at N/P 12. TPGS-LNP produced consistently higher expression than the other formulations across all ratios, with higest expression at N/P 16. Importantly, no apparent cytotoxicity was observed after 24 hours of incubation with any of the three LNP formulations at these N/P ratios (Figure 1e). Based on the balance of efficacy and safety, an N/P ratio of 12 was selected for all subsequent experiments. At this ratio, all three LNP types showed high mRNA encapsulation efficiency ranging from 80 to 90% (Figure 1f).

To assess the stability of the encapsulated mRNA, LNPs-encapsulated mCherry mRNA and non-encapsulated mCherry mRNA were incubated in the kidney perfusion solution (RBC-free) at 37°C for up to 24 hours, with samples collected at multiple time points. The remaining mCherry mRNA levels were quantified by RT-qPCR. Free mRNA exhibited rapid degradation, with a sharp decline in detectable levels within the first hour of incubation. In comparison, mRNA encapsulated within all three LNP formulations remained stable over 24 hours (Figure 1g).

The transfection efficiency of F127-, TPGS- and DSPE-LNP for mRNA delivery was assessed in vitro by quantifying uptake of translationally inactive Cy5-labeled mRNA encoding EGFP (Cy5-EGFP) in OK cells by flow cytometry and by measuring the expression of mRNA encoding EGFP (EGFP) in HKC-8 cells using flow cytometry. Both mRNA uptake (Cy5 signal) and protein expression (EGFP signal) were quantified using flow cytometry. No significant difference in uptake was observed among the three formulations at 4 hours, while DSPE-LNP exhibited the highest cellular uptake after 24 hours (Figure 2a,b). Regarding mRNA-mediated protein expression, TPGS-LNP demonstrated the highest EGFP levels at 24 hours (Figure 2 a,c). Lipofectamine 2000 (LIPO) was used as a positive control and exhibited higher mRNA uptake while lower reporter protein expression compared with all three LNP formulations.

**Figure 2:**
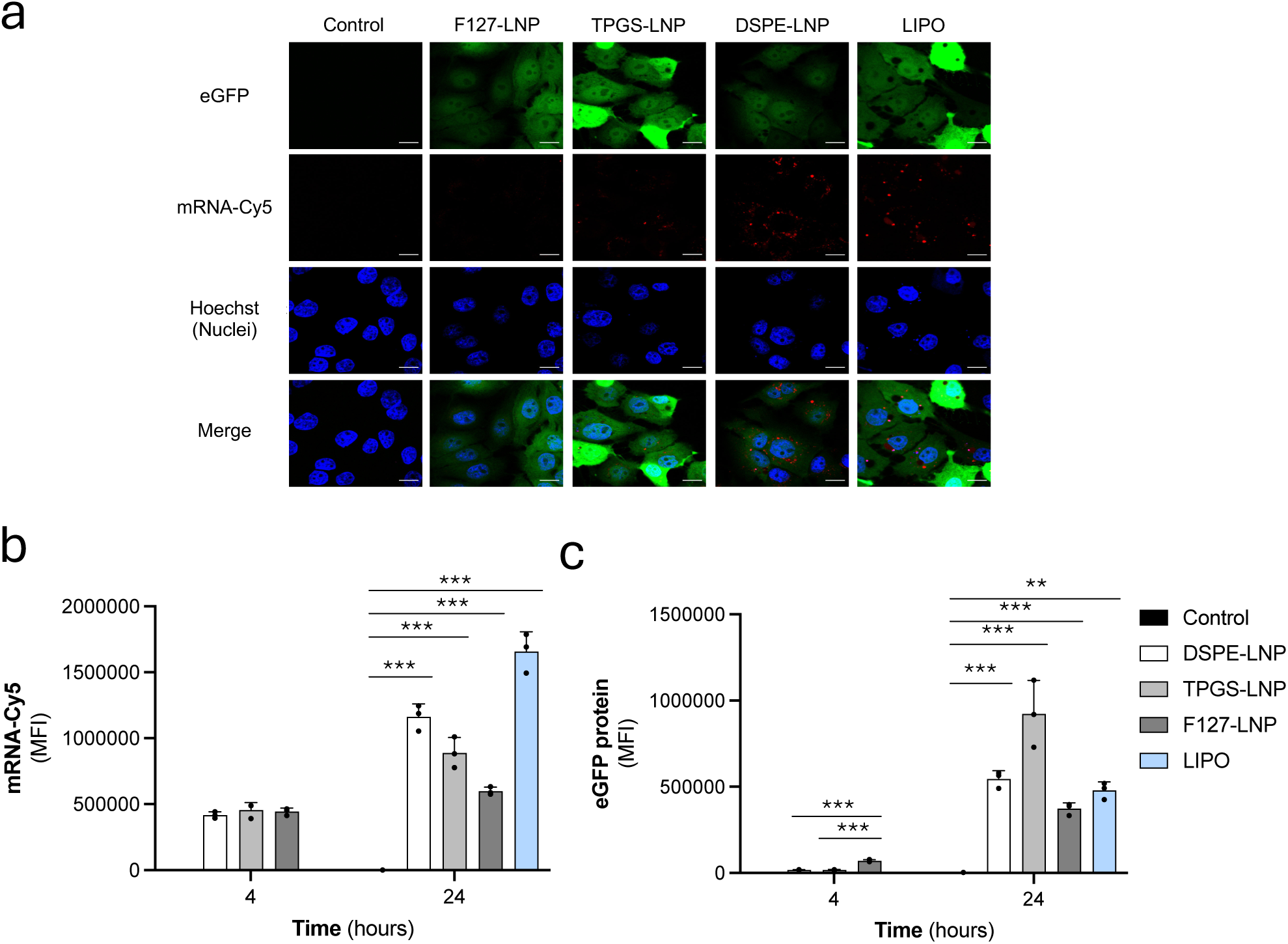
mRNA uptake and protein expression in OK cells transfected with DSPE-LNP, TPGS-LNP and F127-LNP. (a) Representative confocal images of OK cells transfected with LNP encapsulated with Cy5-labeled eGFP mRNA. Green: eGFP protein expression; red: Cy5-labeled eGFP mRNA (mRNA-Cy5); blue: nucleus stained with Hoechst 33342. (b) Quantitative analysis of cellular uptake of Cy-5 labeled eGFP mRNA and (c) eGFP protein expression. Data are presented as Mean ± SD (n = 3). Statistical significance was determined by multiple mixed effects analysis.* p < 0.05, ** p < 0.01, *** p < 0.001.

To further evaluate secreted protein expression in a non-invasive manner, we selected human erythropoietin (EPO) as a model reporter, as its secretion allows quantification in perfusate and urine. As an initial assessment, OK cells were transfected with hEPO mRNA encapsulated in LNPs, and tested in both cell lysate and cell culture supernatant. (Supplemental figure 2). Similar trends were observed using both flow cytometry and ELISA with these alternative reporter proteins, with TPGS-LNP consistently being the most efficiently expressing candidate.

### hEPO mRNA delivery and translation during kidney-NMP

To evaluate mRNA delivery and translation during NMP, human erythropoietin (hEPO) encoding mRNA was chosen as a reporter, enabling direct measurement of secreted hEPO in both the perfusate and urine. Porcine kidneys were connected to an NMP setup and a 5 mL solution containg 150 µg of DSPE (n=4) or TPGS (n=3) encapsulated hEPO mRNA (DSPE-hEPO and TPGS-hEPO, respectively) was infused intra-arterially at 120 mL/h. The contralateral kidneys (n=6) served as non-mRNA treated controls. After LNP-mRNA infusion, kidneys remained on NMP for an additional six hours for follow-up. hEPO was measured in the perfusate as early as 2 hours following mRNA administration, with levels gradually increasing to about 4 mIU/mL for DSPE-hEPO after 6 hours. TPGS-hEPO treated kidneys showed significantly higher hEPO levels with up to 360 mIU/mL after 6 hours (Figure 3a). hEPO was also detected in the urine reaching up to 6 mIU/mL and 420 mIU/mL for DSPE-hEPO and TPGS-hEPO, respectively, measured between 4 and 6 hours after administration depending on individual kidney urine output (Figure 3b).

**Figure 3:**
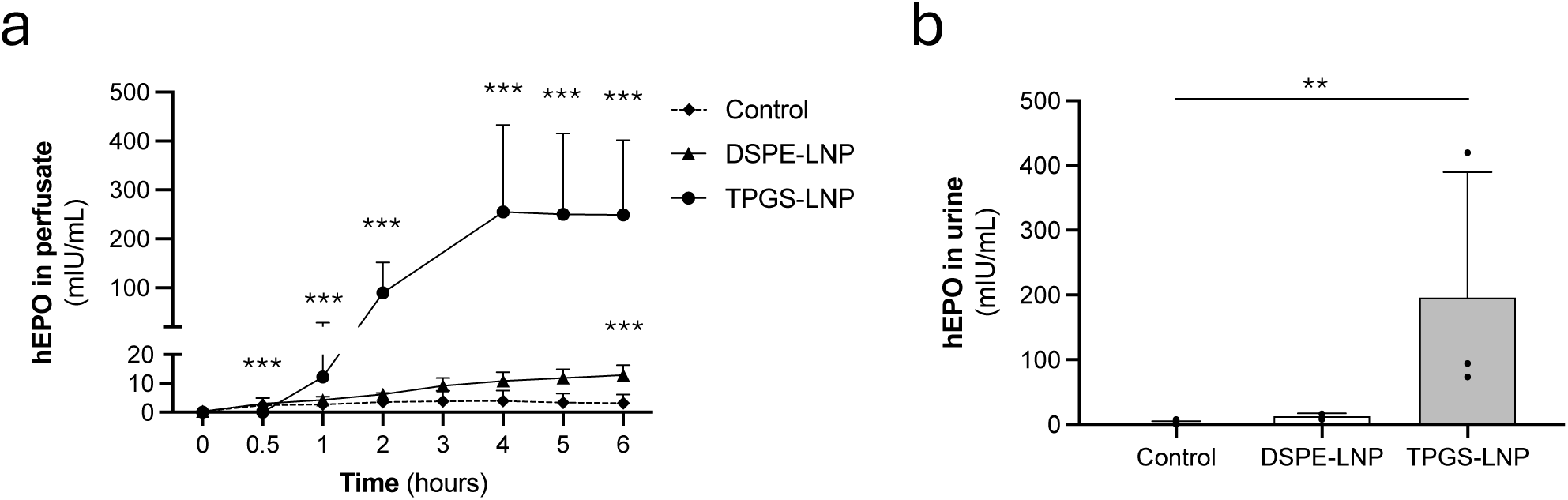
Levels of human erythropoietin (hEPO) detected in perfusate (kidney perfusion solution) and urine following arterial infusion of 150 μg hEPO mRNA via DSPE- and TPGS-LNP during NMP of porcine kidneys. hEPO levels detected by ELISA. (a) Human EPO levels measured in the perfusate over 6 hours. Significance of TPGS-LNP or DSPE-LNP against the control is determined by 2-way ANOVA test with Šídák’s multiple comparisons. * p < 0.05, ** p < 0.01, *** p < 0.001. (b) Human EPO levels detected in the urine 4-6 hours after mRNA infusion. Significance is determined by Kruskal-Wallis. * p < 0.05, ** p < 0.01, *** p < 0.001.

Perfusion characteristics, including renal blood flow, oxygen consumption and urine production, were similar between porcine kidneys infused with DSPE-hEPO, TPGS-hEPO and non-treated controls (Supplemental Figure 3). Hematoxylin and eosin (H&E) as well as periodic acid-schiff (PAS) staining of the tissue sections showed no signs of LNP-induced structural damage (Supplemental Figure 4).

### mCherry mRNA delivery and translation during kidney-NMP

To asses the possibility for localized expression of an intracellular protein, mRNA encoding mCherry was administered to porcine kidneys using both DSPE- and TPGS-LNP, following the same protocol as in the hEPO studies. Pilot studies showed initial mCherry protein expression around 6 hours, therefore perfusion time was extended to 9 hours.

In tissue biopsies taken hourly up to 9 hours of NMP, western blotting showed apparent mCherry protein expression after 4-5 hours for TPGS-LNP and 7-8 hours for DSPE-LNP, with protein levels increasing progressively toward the end of the perfusion period (9 hours; Figure 4a, b). Consistent with the findings observed for hEPO expression, TPGS-LNPs exhibited greater overall efficacy compared with DSPE-LNPs (Figure 4a, b). Analysis of cortical samples obtained from both kidney poles, as well as medullary tissue, confirmed that the delivered mRNA reached all anatomical regions of the organ. Nevertheless, substantial inter-kidney variability in mCherry protein levels was observed.

**Figure 4:**
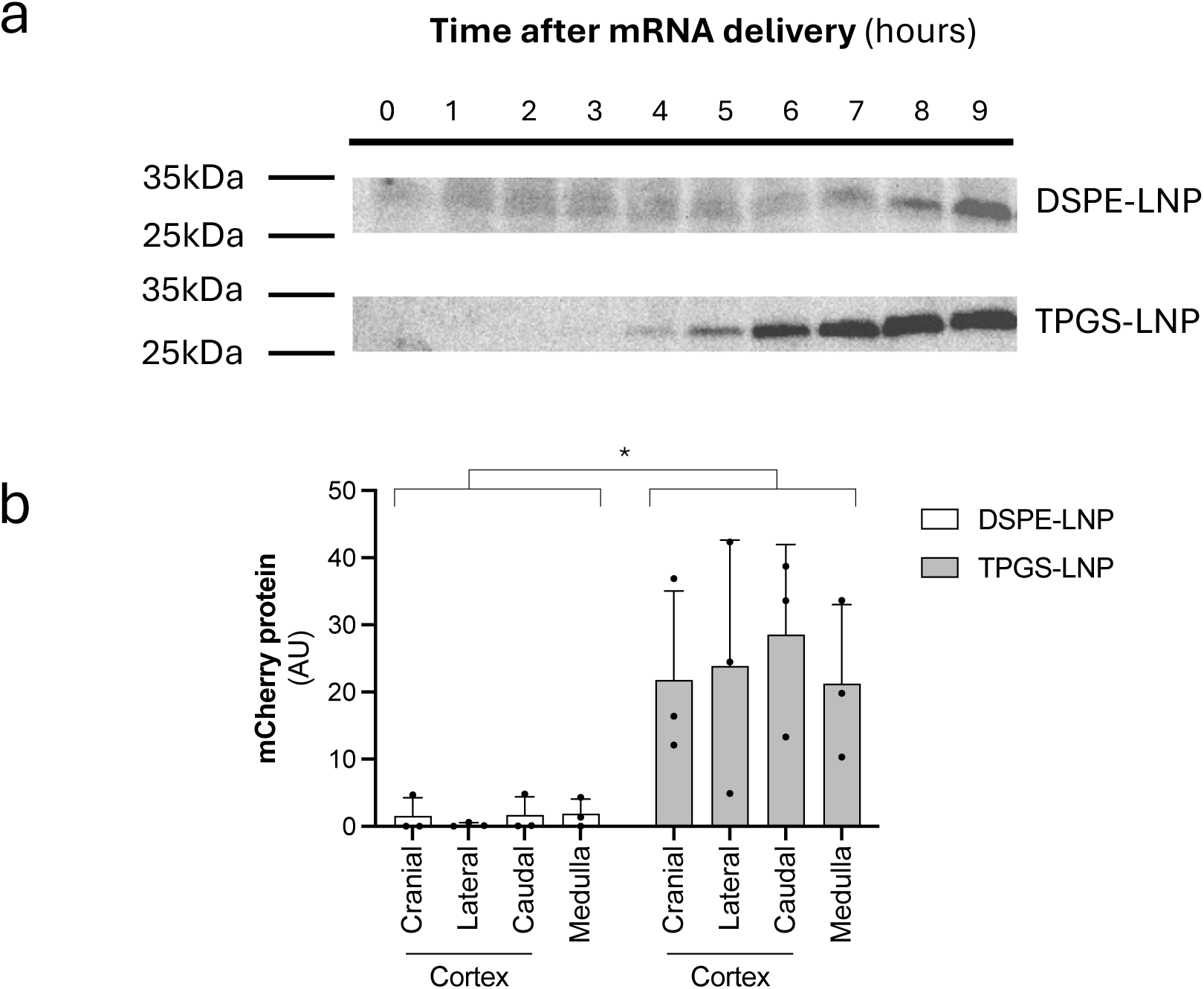
Levels of mCherry detected by western blot throughout several kidney regions following arterial infusion of 150 μg mCherry mRNA via DSPE- and TPGS-LNP during NMP of porcine kidneys. (a) Representative western blot showing mCherry expression (28 kDa) in kidney cortical tissue after delivery with DSPE- and TPGS-LNPs. (b) Bar plot quantifying mCherry levels in different kidney regions, including the cortex (cranial, lateral, caudal) and medulla obtained after the end of the perfusion Signals were quantified using ImageLab software and normalized to total protein signal from the corresponding stain-free gel. Significance is determined by two-way ANOVA with Šídák’s multiple comparisons. * p < 0.05, ** p < 0.01, *** p < 0.001.

During the 9-hour follow-up period, high intrinsic autofluorescence in porcine kidney tissue precluded reliable detection of native mCherry fluorescence, limiting direct fluorescence-based assessment of protein expression. Therefore, immunohistochemistry (IHC) was used to evaluate mCherry protein localization at the cellular level. IHC revealed specific mCherry immunoreactivity in kidneys treated with mRNA-LNPs, whereas untreated control kidneys showed no detectable signal (Figure 5a, c, e). mCherry protein expression was predominantly localized to scattered tubular epithelial cells and displayed a heterogeneous, patchy distribution across the renal parenchyma. Glomerular and vascular structures showed minimal to no detectable mCherry protein staining.

**Figure 5:**
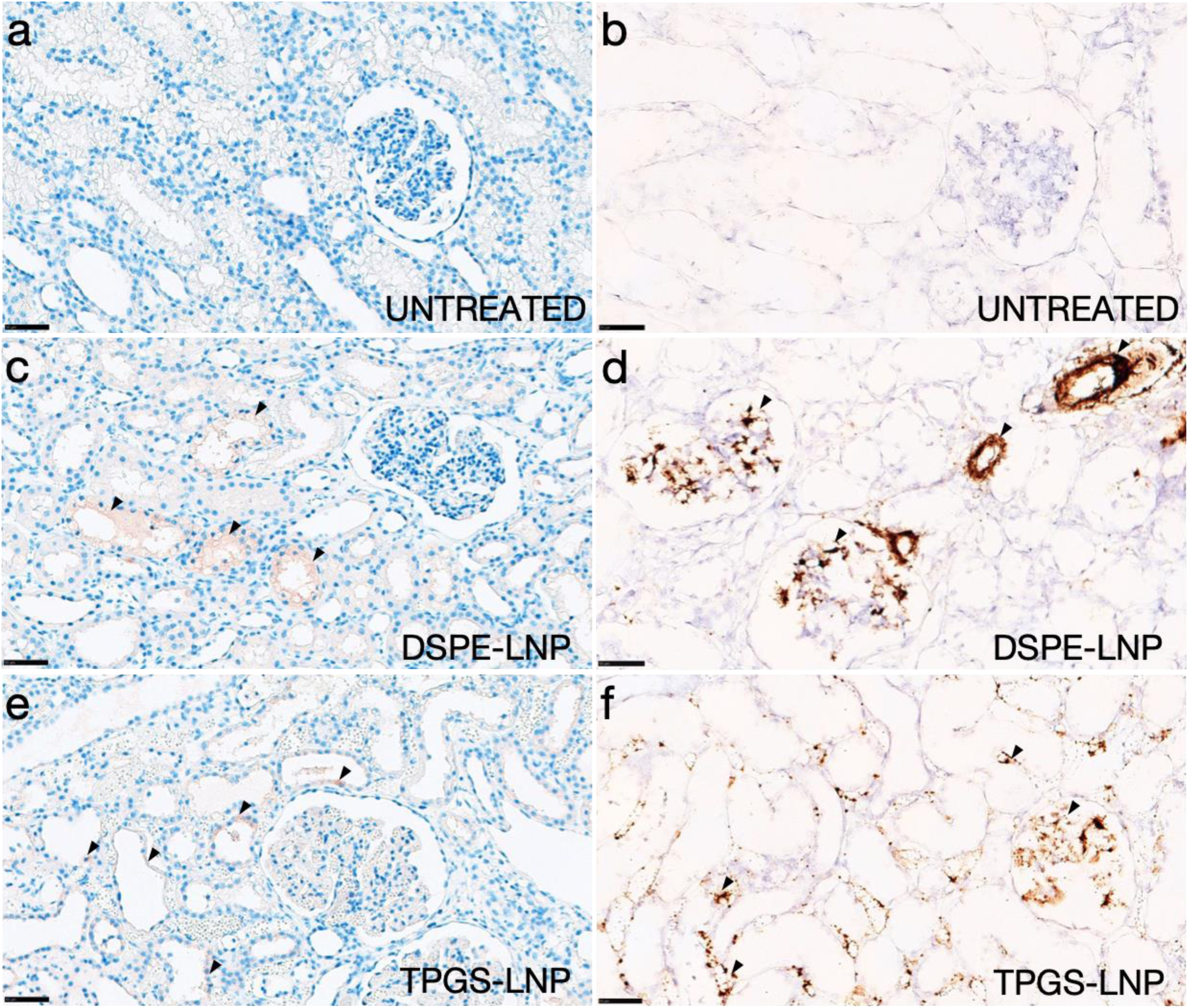
mCherry mRNA and mCherry protein presence in kidney tissue following arterial infusion of 150 μg mCherry mRNA via DSPE- and TPGS-LNP during NMP of porcine kidneys. mCherry protein was detected by immunohistochemistry (IHC), and mCherry mRNA was detected by brightfield RNA in situ hybridization (BRISH). (A, C, E) Representative brightfield images of mCherry protein expression visualized by IHC using a brown DAB chromogen. (B, D, F) Representative brightfield images of mCherry mRNA detected by BRISH with hematoxylin counterstaining. Panels A and B show tissue from an untreated control kidney perfused under identical conditions without mRNA-LNP administration, panels C and D show tissue from a kidney treated with TPGS-LNPs, and panels E and F show tissue from a kidney treated with DSPE-LNPs. Positive mCherry signal is indicated by arrowheads. Images are shown at high magnification. Scale bars represent 50 µm.

To further characterize the intrarenal distribution of the delivered transcript, branched RNA in situ hybridization (BRISH) was performed to detect mCherry mRNA. BRISH analysis demonstrated widespread intracellular mCherry mRNA signal throughout the tissue in treated kidneys, whereas untreated control kidneys showed no detectable signal, highlighting the effective tissue penetration of both LNPs after *ex vivo* intra-arterial administration (Figure 5b, d, f). mCherry mRNA was detected in multiple renal compartments, including glomeruli, tubular epithelial cells, and vascular-rich regions, consistent with effective tissue penetration following intra-arterial delivery during normothermic machine perfusion.

## DISCUSSION

This study establishes a clinically compatible strategy for organ-directed mRNA delivery during *ex vivo* perfusion. By delivering mRNA to the isolated organ, off-target effects commonly associated with systemic mRNA delivery are minimized, including hepatic interactions (9, 10, 21). Intra-vascular LNP administration via the renal artery achieved effective tissue penetration and biodistribution of the reporter mRNAs. hEPO expression became detectable approximately 2 hours after administration in both perfusate and urine, while mCherry expression was observed in the renal parenchyma after 6 hours.

Building upon previously validated ionizable lipid-based LNP formulations (20, 22), we evaluated three different PEG-lipids: Pluronic F127, TPGS, and DSPE-PEG2000, each contributing unique properties (23–26). While all formulations showed comparable physical characteristics, DSPE-PEG2000 and F127 exhibited slightly higher mRNA encapsulation efficiencies than TPGS, potentially due to TPGS’s smaller lipid head group and shorter PEG chain, which may compromise particle stability.

In spite of this, our results indicate that during ex vivo delivery to whole porcine kidneys, TPGS-containing LNPs achieved markedly higher protein expression compared to the conventional DSPE-PEG2000. This correlates with the *in vitro* findings, where TPGS-LNP outperformed the other formulations in terms of expression of the reporter protein. The enhanced potency of TPGS-containing LNPs under organ-confined *ex vivo* conditions may be partly explained by the ability of TPGS to inhibit P-glycoprotein activity(24, 27, 28) which is highly expressed in proximal tubule cells and plays a central role in efflux transport. Additionally, the improved solubility and permeability conferred by TPGS may enhance biocompatibility and tissue accumulation, further enhancing delivery efficiency(29).

Our previous *in vivo* work in rodents using systemic administration reported effective dosing from 0.01 to 1 mg mRNA/kg(22), while clinical studies employing local mRNA delivery-such as epicardial injection or nebulized pulmonary administration-have used substantially higher absolute doses in the 3–24 mg mRNA/kg range (5, 11). Although *ex vivo* perfusion differs fundamentally from *in vivo* delivery, these data was used as a general benchmark for establishing a starting dose. Therefore, based on these and previous works (30–32), we selected a dose of 0.15 mg mRNA per porcine kidney to assess feasibility in the *ex vivo* organ perfusion setting. Notably, the relatively low dose of 0.15 mg mRNA per porcine kidney (weighing approximately 100–150 g) combined with the intra-arterial organ-confined administration, resulted in high levels of the reporter protein human EPO reaching up to 400 mIU/mL in the perfusate. For comparison, normal circulating EPO concentrations in healthy humans range from 2.8 to 17.1 mIU/mL(33).

*Ex vivo* mRNA delivery was successfully achieved using human-sized porcine kidneys. Furthermore, LNPs were administered through a simple infusion via the arterial port of the NMP setup, confirming the clinical feasibility of the approach.

LNP-mRNA treatment was well tolerated during NMP, with preserved perfusion parameters and renal architecture. No evidence of cytotoxicity was observed *in vitro* and *ex vivo*. Importantly, the potential adverse effects associated with systemically delivered LNP-mRNA studies are not anticipated in our ex vivo approach, as systemic exposure is minimized. Moreover, following NMP, it is standard clinical procedure to flush the kidney and/or return it to hypothermic machine perfusion prior to transplantation, which facilitates the removal of any free LNPs.

The rapid appearance of the mRNA provide a window in which the encoded protein may become available during the critical post-transplant period, potentially driving new therapeutic strategies to mitigate ischemia reperfusion injury (IRI), promote tissue regeneration, and transiently modify the donor organ during preservation, enhancing graft resilience during the critical period immediately following transplantation(15, 17, 34). Given the complex and multifactorial nature of IRI, it may be necessary to target multiple pathways(17). Donor kidneys would greatly benefit from a combined and individualized treatment strategy and mRNA therapy offers a flexible platform for such an approach, as it can encode both gain-of-function activators and inhibitors, which could potentially be combined into a single solution. Assessing post-transplant benefit in transplant models is a logical next step but beyond the present ex vivo feasibility study.

In summary, we have successfully developed and screened LNPs for efficient kidney-confined mRNA delivery and on-demand protein expression during NMP and potentially after implantation. TPGS-based LNPs has emerged as a promising vector for mRNA delivery and expression to an isolated kidney, demonstrating their efficacy both *in vitro* and *ex vivo* in human-sized porcine kidneys. Furthermore, we established a practical strategy for using mRNA therapeutics in donor organs, providing a foundation for their application in *ex vivo* NMP as a precision medicine platform prior to organ transplantation.

## METHODS

### Cell culture

Opossum kidney cells (OK cells) were cultured in low glucose Dulbecco’s Modified Eagle Medium (DMEM) (Thermo Fisher, Waltham, MA), supplemented with 10% fetal bovine serum (FBS, Thermo Fisher, Waltham, MA) and 1% penicillin-streptomycin (Thermo Fisher, Waltham, MA) at 37°C, 5% CO_2_ and 100% humidity.

### Preparation of nanoparticles

Lipid nanoparticles (LNP) were synthesized through nanoprecipitation. Briefly, Pluronic (F127, Sigma-Aldrich), C14120(35), cholesterol (Sigma-Aldrich), 1,2-dioleoyl-sn-glycero-3-phosphocholine (DOPC, Avanti Polar Lipids) and d-α-tocopheryl polyethyleneglycol 1000 succinate (TPGS, Sigma-Aldrich) were dissolved in 100% ethanol. For F127-LNP, F127 and C14120 were combined at the weight ratio of 2:1. For DSPE-LNP and TPGS-LNP, C14120, cholesterol, DOPC, and the PEG-Lipid either TPGS or 2-distearoyl-sn-glycero-3-phosphoethanolamine-N-[methoxy(polyethylene glycol)-2000] (DSPE-PEG2000, Avanti Polar Lipids) were combined at weight ratio of 4:2:1:1. Subsequently, sodium acetate (200mM, pH 5.5) was rapidly injected into the organic lipid mixture with vigorous stirring, leading to be self-assembly of empty nanoparticles. Ethanol was removed by dialysis against 200 mM sodium acetate pH 5.5 using a 3.5 kDa dialysis tube (Thermo Scientific). LNPs were stored at 4°C before further experiments.

### mRNA encapsulation and delivery

The following mRNAs were used: CleanCap EPO mRNA (5moU, Trilink, L-7209), CleanCap mCherry mRNA (5moU, Trilink, L-7203) and CleanCap Cy5 eGFP mRNA (5 moU, Trilink, L-7701). mRNA was encapsulated through electrostatic interaction when thoroughly mixed with the LNPs.

For in vitro studies, LNPs (1 mg/mL, concentration of C14120) in 200 mM sodium acetate buffer pH 5.5) were mixed with mRNA (at 200 ng/μL) at different N/P ratios. 100□ng of mRNA was mixed with  □0.75□µL, 1.5□µL, and 2□µL of LNPs to obtain N/P ratios of 6, 12 and 16, respectively. The LNP/mRNA mixture was incubated for 10 minutes at room temperature (RT) and added to the culture medium.

For delivery during kidney NMP, 1100 μg LNPs (in 2.5 mL 200 mM acetate buffer, pH 5.5) were mixed with 150 μg of mRNA (prediluted in 0.5 mL of nuclease-free water) and incubated for 10 minutes at RT. Prior to the infusion, 2 mL of isotonic saline (0.9 mg/mL NaCl Fresenius Kabi, B05BB01) was added. This solution was transferred to a 20 mL syringe and administered via the arterial side of the NMP circuit at an infusion rate of 120 mL/h, starting 30 minutes after the initiation of kidney-NMP. Following the initial infusion, the line was flushed with 3 mL of isotonic saline.

### Characterization of LNPs

The hydrodynamic diameter and surface charge of the LNPs were determined by photon correlation spectroscopy (PCS) and Laser Doppler Velocimetry (LDV) at 25 °C on a Zetasizer Nano ZS (Malvern Instruments, Malvern, UK). The encapsulation efficiency of mRNA was quantified using the RiboGreen assay (Invitrogen, Copenhagen) following the manufacturer’s protocol. Briefly, 50 μL of LNP/mRNA solution was mixed with 50 μL of RiboGreen working solution (1:100 in TE buffer: 10 mM Tris, 0.1 mM EDTA, pH 7.5). After 5 minutes of incubation at RT, the fluorescence intensity was measured by FLUOstar OPTIMA (Moritex BioSience) at an excitation/emission wavelength of 480/520 nm.

### Cell viability test

Cell viability was evaluated via AlarmaBlue (Molecular Probes, Life Technologies) assay according to the instructions from the manufacturer. OK cells were seeded in a 96-well plate at a density of 1×10^4^ cells/well. The following day, the growth medium was refreshed and each well was transfected with 33 ng LNP-encapsulated mCherry mRNA. After 24 hours, growth medium supplemented with 10% (v/v) of AlarmaBlue reagent was added. After 2 hours of incubation, 80 μL of the supernatant was transferred to an opaque 96-well NUNC plate (Sigma-Aldrich, P8616), and the fluorescence intensity was measured using a FLUOstar OPTIMA plate reader (Moritex BioScience) at excitation and emission wavelengths of 540 nm and 590 nm, respectively.

### Stability of mRNA encapsulated in LNPs

To assess stability, 88 ng of LNP-encapsulated mCherry mRNA was incubated in 45 μL of NMP perfusate (see supplemental table 1) at 37°C for 0 minutes (right after mixing), 15 minutes, 30 minutes, 1 hour, 2 hours, 4 hours, 6 hours and 24 hours. After incubation, lysis buffer (0.25% Igepal CA-630, 10mM Tris·HCl pH 7.4, 150 mM NaCl, and 1 U/μL RNase inhibitor) was added and samples were centrifuged at 10,000 g for 5 minutes. Next, 10 μL of supernatant was used directly for cDNA preparation using Revert Aid RT Reverse Transcription Kit (Thermo Fisher). The cDNA was diluted 40 times, and 4 μL was used as template for quantitative real-time PCR using SYBR Green Master Mix (Thermo Fisher), running on a Light Cycler 480 Real-Time PCR System (Roche). The primer sequences for mCherry are: Forward 5′-CGGGTGATGAACTTCGAGGA-3′ and Reverse 5′-TCGAAGTTCATCACCCGCTC-3′.

### In vitro performance of LNPs for mRNA delivery

OK cells were seeded in 24-well plate (1×10^5^ cells/well). The next day, growth medium was refreshed and 200 ng LNP-encapsulated mRNA was added per well. After 24 hours, cells were detached using 0.05% Trypsin-EDTA (Thermo Fisher, Waltham, MA) and fluorescent intensity was quantified by flow cytometry (NovoCyte, ACEA Biosciences).

For microscopy studies, OK cells were seeded in 8-well µ-Slide (ibiTreat, ibidi; 5×10^4^ cells/well). The next day, the growth medium was refreshed and 100 ng LNP-encapsulated mRNA was added per well. After 24 hours, medium was replaced by FluoroBrite DMEM (Thermo Fisher Scientific, Waltham, MA), and cells were imaged by confocal laser scanning microscopy (LSM 710, Zeiss, Germany) and analyzed by Fiji ImageJ software 2.3.0.

### Organ procurement and normothermic machine perfusion

A comprehensive account of the ethical approval process, surgical techniques and blood processing procedures is available in Supplemental Material 1.

Porcine kidneys were procured under general anesthesia following a standardized surgical protocol, including midline laparotomy and bilateral nephrectomy with a warm ischemia time as in a living donation procedure. Special care was taken to preserve vascular integrity and obtain sufficient arterial length for optimal cannulation. The renal artery was cannulated with a straight t-cannula immediately post-nephrectomy and with minimal warm ischemic time. Then, the kidney was flushed with heparinized saline followed by cold preservation solution before being stored at 4 °C. Further procedural details are provided in the Supplemental Material 1.

After a maximum of 25 hours of cold ischemia, all kidneys were connected to an open clinically compatible NMP system via the arterial cannula and venous outflow drained freely into the reservoir under gravity, permitted via an uncannulated renal vein.

The NMP circuit was primed with 600 mL human albumin erythrocyte-based solution (see composition supplemental material 2). The perfusate temperature was kept at 37°C and oxygenated with 0.5 L/minute carbogen gas (95% O2, 5% CO2). After the kidney was connected to the system, a mean arterial pressure of 80 mmHg was maintained. Gas analysis was carried out every hour (ABL90 FLEX, Radiometer) and hourly perfusate, urine and biopsies were taken. Urine output was measured, and the rest of the urine was recirculated. During NMP, glucose and pH were adjusted if glucose dropped under 5 mmol and pH under 7.35 by adding a bolus of glucose 5% (50 mg/mL, Fresenius Kabi, 7340022100429) or sodium bicarbonate 8.4% (8.4 g/mL, Fresenius Kabi, 03058282) directly to the organ chamber.

### Sampling and storage

During follow-up on NMP, perfusate from the vein and urine samples were collected. Samples were centrifuged for 12 minutes at 3000 g at 4°C and supernatant was stored at −80°C. Blood gases from arterial and venous perfusate as well as urine, were measured hourly (ABL90 FLEX, Radiometer) to monitor the perfusion process. Each 6 mm biopsy was stored in RNA-later (Merck KGaA) at −80°C. At the end of the perfusion 1×1 cm biopsies with both cortex and medulla were taken and formalin fixed (Biopsafe, 3178-200-03) for about 24 hours at 4°C, and were then changed to phosphate buffered saline until embedding.

### Human EPO ELISA

Human EPO protein expression in kidney NMP perfusate and urine samples was quantified by a Human EPO ELISA (Invitrogen, BMS2035-2). Human EPO in OK cell cultures was quantified by an EPO DuoSet ELISA (R&D Systems, DY286-05) according to the manufacturer’s protocols.

### mCherry Western Blot

In total, 25 mg of kidney tissue was homogenized in M tubes (Miltenyi Biotec,130093236) in 500 µL of Pierce RIPA buffer (Thermo Scientific, 89901) supplemented with cOmplete Mini protease inhibitor cocktail (Merck KGaA, 11836153001). Lysates were centrifuged (10 minutes, 3000 g, 4°C) and supernatant was stored at −80°C until further use. Total protein was quantified using Pierce BCA Protein Assay Kit (Thermo Fisher, 23225) according to the manufacturer’s protocols. In total, 30 µg of protein was run on an Any kD CriterionTM TGX Stain-Free gel (BIO-RAD, 5678124) and transferred onto nitrocellulose Trans-Blot®TurboTM membranes (BIO-RAD, 1704159). Blots were blocked using 5% Skim milk powder (Millipore, 999999-99-4) and washed in 20 mM TBS-T (Tris-buffered saline, pH 7.6, 0.1% TWEEN 20). Membranes were incubated with 0.35 μg/ml DSHB-mCherry-3A11-s antibody (Development Studies Hybridoma Bank, DSHB) in 20 mL of TBS-T overnight at 4°C, followed by 1 hour of incubation at RT. After being washed in TBS-T, blots were incubated with 0.4 µg/ml polyclonal Goat Anti-mouse Immunoglobulins/HRP antibody (Aligent, P0447) in TBS-T for 1 hour at RT. After wash, blots were incubated with 3 mL ECLTM Prime Western Blotting Detection Reagent (Cytvia, RPN2236) for 5 minutes at RT, and imaged using the ChemiDocTM MP Imaging System (BIO-RAD). Quantification of bands in the blot was done with ImageLab (version 6.0.1, 2017) and signal was corrected for total protein of the respective gel.

### Immunohistochemistry (IHC) and microscopy

The formalin-fixed paraffin-embedded (FFPE) samples were sectioned at 2.5 µm with a microtome (Thermo Scientific, HM3555). Hematoxylin and eosin (H&E) and Periodic Acid–Schiff (PAS) staining were performed on FFPE kidney sections (3–4 µm thick) using standard protocols. H&E was used to evaluate general tissue morphology, while PAS staining was used to assess glomerular basement membranes, tubular brush borders, and mesangial matrix integrity. Histological scoring was performed by a trained pathologist blinded to the treatment groups. Kidney sections were evaluated for features including tubular damage, dilatation, cast formation, vacuolization, glomerular injury, interstitial inflammation, and edema. Each parameter was semi-quantitatively scored based on predefined criteria, and overall tissue integrity was assessed to compare damage and structural preservation between groups.

Immunohistochemistry (IHC) was performed on FFPE porcine kidney sections using a Ventana Discovery Ultra automated staining platform (Ventana Medical Systems). Sections were deparaffinized and rehydrated using the manufacturer’s standard automated protocols. Antigen retrieval was performed using CC2 buffer (Ventana Medical Systems) at 99 °C for 30 min. Endogenous peroxidase activity was blocked prior to primary antibody incubation. A rabbit monoclonal antibody against mCherry (EPR20579; Abcam; RRID: AB_213511) was applied at a dilution of 1:100 and incubated for 32 min at 36 °C. Signal amplification was achieved using DISCOVERY OmniMap anti-rabbit HRP (Ventana Medical Systems). Visualization was performed using the OptiView DAB IHC Detection Kit (760-700; Ventana Medical Systems), followed by hematoxylin counterstaining. Negative controls consisted of kidneys perfused under identical conditions but without mRNA-LNP administration. Whole-slide images were acquired using a Hamamatsu NanoZoomer XR digital slide scanner. Digital image analysis and visualization were performed using NDPview2 software (version 2.9.29; Hamamatsu, macOS).

### Brightfield RNA In Situ Hybridization (BRISH)

RNA-ISH was performed by using the RNAscope Probe-mCherry-O10-C1 (Cat. No. 1264111-C1, target region 2-714 bp), which is a probe specific mCherry mRNA. For RNA quality controls, probes targeting the housekeeping gene peptidylprolyl isomerase B (cyclophilin B) were used as follows: Ss-PPIB (Cat. No. 428591, target region 16-642 bp) for pig tissue, and Hs-PPIB (Cat. No. probe. 313901, target region 139-989 bp) for cells.

As specificity control for mCherry detection, about 100.000 HEK293 cells were transfected with 200 ng mCherry mRNA carried by 1.5 μg of TPGS-LNP for 24 hours. HEK293 cells incubated for 24 hours without transfection were used as controls. Cells were fixed with 4% paraformaldehyde (PFA) for 1 hour and placed in agar solution (0.18 g of agar in 10 mL of demineralized water). Followed by a second fixation with 4% PFA and paraffin embedding.

RNA-BRISH was performed on 5-µm-thick FFPE sections on SuperFrost PLUS microscope slides. Pretreatment was done following the RNAscope Sample Preparation and Pretreatment Guide for FFPE Tissue (Cat. No. 322381). This included blocking H_2_O_2_ by adding the RNAscope hydrogen peroxide to the slides and incubating them for 10 minutes at RT. Antigen retrieval was performed by heating the slides for 18 minutes in the RNAscope antigen retrieval buffer (1:10 dilution and 98-100 °C) using a steamer. Then, protease treatment was performed by adding the RNAscope Protease Plus to the sections and incubating them at 40°C for 30 minutes in the HybEZ oven, followed by probe hybridization overnight. Signals were detected by using the RNAscope 2.0 HD Detection Kit (Cat. No. 322310). This included amplification by a series of amplification steps and signal detection by using DAB (3,3’-Diaminobenzidine). The reactivity was visualized as brown dots. Following BRISH, nuclear counterstaining was performed using hematoxylin. Sections were coverslipped and digitized using an Olympus SLIDEVIEW VS200 slide scanner (Olympus Nederland B.V., Hoofddorp, Netherlands).

### Data analysis

GraphPad Prism (version 10.2.2) was used to perform statistical analyses and compose graphs. Data are presented as mean ± SD unless otherwise stated. Statistical tests included two-way ANOVA, t-tests, and appropriate non-parametric tests based on data distribution. Statistical significance was set to p < 0.05.

## DATA AVAILABILITY STATEMENT

Raw numerical data underlying Figures 1-4 and Supplementary Figures S1-S4 (GraphPad Prism files) are available upon request.

## ACKNOWLEDGEMENTS

The authors would like to express their gratitude to Mirjiam Mastik for her valuable assistance in carrying out the BRISH studies. Scoring of kidney tissue sections was supported by pathologist Søren Rasmus Palmelund Krag. And lastly, the authors would like to acknowledge Lene Mejer Rosenlund, Maria Arge and Birgitte Kildevæd Sahl for their expert biotechnician assitance. This work was supported by Augustinus Fonden and Novo Nordisk Foundation grant NNF23OC0081177 (RNA-META).

## AUTHOR CONTRIBUTIONS

ME, JK, BJ, AK, RN, HL, CBB and PS conceptualized the research design. CBB, PS, KBK, VRL, CJ and AVF participated in the performance of the experiments. CBB, PS, HH and JH participated in the analysis. CBB and PS drafted the manuscript. ME performed critical revision. ME, JK, BJ and AK supervised the project and reviewed the final version of this manuscript. All authors approved the final version of the manuscript.

