## Supplementary for "Organ-confined mRNA delivery and protein expression during *ex vivo* perfusion of porcine donor kidneys"

**SUPPLEMENTARY MATERIAL 1**

**Ethics**

The study utilized female Danish Landrace and Yorkshire crossbreed laboratory pigs (weight: 50-55 kg) adhering to the European Union (Directive 2010/63/EU). The study was approved by the Animal Experimentation Council in Denmark (reference: 2020-15-0201-00624). All involved personnel were certified by the Federation of European Laboratory Animal Science Associations.

**Surgical procedure**

For the recovery of porcine kidneys, all animals were fasted from midnight on the day of surgery. Animals were sedated prior to transportation to the surgical suite using 0.1 mL/Kg of “Zoletil-mix” (Zoletil 50 vet. (125 mg Tiletamine, 125 mg Zolazepam), 125 mg Xylazine, 125 mg Ketamine, 25 mg Butorphanol) intramuscularly (IM). Upon arrival, a 18G intravenous (IV) catheter was placed in the ear. Intubation was performed using a 7.5 mm endotracheal tube, which was secured to the snout of the pig to avoid displacement. The animal was then connected to the respirator and ventilated using tidal volumes of 4-6 mL/Kg at a fractional inspiratory O2 of 30-50%, aiming at a saturation > 98% and an end tidal CO2 between 4.5-5.5 KPa. Blood pressure and heart rate were monitored, maintaining a heart rate between 45-100/minute and a systolic blood pressure > 100 mmHg. After induction, general anesthesia was maintained by 4-8 mg/Kg/h of propofol (10 mg/mL, Baxter A/S, 458982) and analgesia was accomplished with 6-10 µg/Kg/h fentanyl (50 µg/mL, B Braun Medical, 071036). Temperature was measured using a rectal probe. The external jugular vein was cannulated with an 8Fr, 10 cm central venous catheter (CVC, Radifocus, Terumo Corporation, RS-B80N10MRD) under ultrasound guidance for fluid administration with isotonic saline (NaCl 0.9%, Fresenius Kabi, B05BB01) and posterior blood collection. Surgery commenced with an approximately 20-25 cm midline laparotomy and kidneys were exposed retroperitoneally using a no-touch technique. Both kidneys were prepared for removal and a papaverine and isotonic saline mixture (1 mL of papaverine 20 mg/mL, 8469245, and 19 mL of NaCl 0.9%, Fresenius Kabi, B05BB01) was injected into the vascular sheath of the renal artery. Both kidneys were clamped using vascular clamps to avoid damage to the vessels and then removed, taking ample length on the artery to ease cannulation during machine perfusion. The ureter was resected distally to the kidney to allow catheterization and posterior urine collection during machine perfusion. After the double nephrectomy, heparin (500 IE/Kg, Panpharma, 482480) was administered to the pig and 5 quadruple-blood bags (Macopharma, LQT613E) of 450 mL of blood were collected. Then, the animals were euthanized with 15 mL sodium pentobarbital (400 ng/mL, Euthanimal, Scanvet A/S, QN51AA01) under general anesthesia.

**Arterial cannulation and storage**

After the nephrectomy of porcine kidneys, the artery of the graft was cannulated with a straight t-cannula (3 or 5 mm, Organ Recovery Systems, CAN0003 or CAN0005) and a cold pre-flush was performed with 20 mL heparinized saline (1 mL of heparin 5000 IE/mL, Panpharma, 482480 in 19 mL saline). Consequently, the kidney was flushed with 300 mL of cold Belzer University of Wisconsin Cold Storage Solution (UW-CSS, Bridge to Life, RM/N 4055). At this point, the ureter was cannulated using either a urosoft ureteral stent set (Angiomed, 57010010) or a 6 Ch feeding tube (ENFit, Medicina Ltd., CG6/120). The kidneys were then preserved at 4ºC on static cold storage until the start of NMP.

**Erythrocyte collection**

Concerning the porcine erythrocytes, following blood collection, blood fractionation (Baxter Optipress II Blood Component Separator, Baxter) was performed, and the leucocyte-filtered erythrocytes were stored in bags containing storage solution saline-adenine-glucose-mannitol at 4ºC (Macopharma, LQT613E). Erythrocytes were then washed in Dulbecco’s phosphate-buffered saline (pH 7.4, Thermo Fisher) between 30 minutes and 1 hour prior to the start of NMP.

**Normothermic machine perfusion**

For the porcine kidneys, NMP circuit consisted of an oxygenator (Maquet Quadrox-i Neonatal, Getinge, 701070415), a centrifugal pump and unit (Medos Deltastream DP2 and Pumpdrive DP2, Medos Medizintechnik AG), a custom-modified LifePort organ chamber (Perfusion circuit, Organ Recovery Systems, LKT200), a custom-made electronic interface and software (LabVIEW Software, National Instruments Netherlands BV) for a pressure-controlled pulsatile perfusion system (100 mmHg systolic pressure and 60 mmHg diastolic pressure, 60 bpm), an ultrasonic clamp-on flow probe (Transonic Systems Europe BV, ME-PXL) to measure flow. Pressure was measured directly with a pressure transducer (TruWave disposable pressure transducer, Edwards Lifesciences). A water bath (Corio CD, Julabo) was used to achieve normothermic temperatures and two syringe pumps (Alaris™ CC Plus) were added to the circuit to the arterial port for additives.

| **Constituent** | **Volume for 1 kidney (mL)** |
| --- | --- |
| **NaCl (9 mg/mL, Fresenius Kabi)** | 200 |
| **Aqua (B Braun)** | 80 |
| **Sodium bicarbonate (8.4 g/mL, B Braun)** | 16 |
| **Calcium gluconate (100 mg/mL, B Braun)** | 4 |
| **Mannitol (150 mg/mL, Fresenius Kabi)** | 5 |
| **Aminoplasmal Elektrolyt (B Braun)** | 5 |
| **Glucose (50 mg/mL, Fresenius Kabi)** | 13 |
| **Magnesium sulphate (200 mg/mL, Region Midt Apoteket)** | 0.5 |
| **Sodium phosphate (150 mg/mL, Region Midt Apoteket)** | 0.15 |
| **Heparin (5000 UI/mL, Panpharma)** | 0.5 |
| **Pen Strep (Gibco, 10000 U/mL)** | 2 |
| **Soluvit multivitamins (Fresenius Kabi, diluted in 10 mL of Aqua)** | 0.5 |
| **Human albumin (200 g/L, CSL Behring)** | 100 |
| **Red Blood Cells** | 250 |
| **Verapamil (0.25 mg/mL, Sigma-Aldrich)** | 10 |
| **Creatinine (Sigma-Aldrich)** | 0.052 g |

Supplemental table 1: Kidney perfusion solution.

**FIGURES**

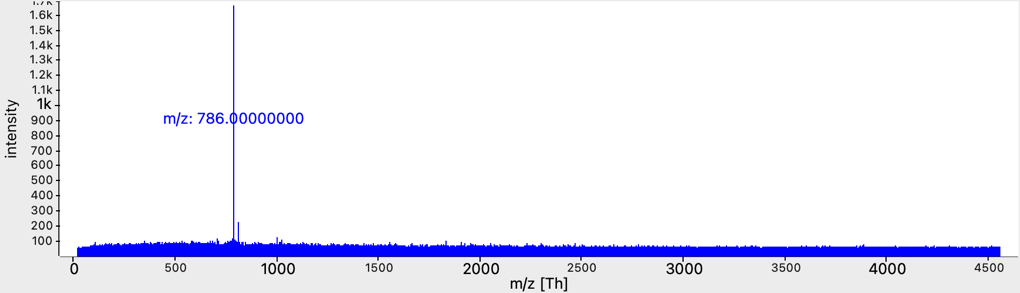

Supplemental figure 1: Structural validation of C14120 by MALDI-TOF mass spectrometry. The observed molecular ion peak ([M+H]⁺) at m/z 787 corresponds to the calculated molecular mass of 786, confirming the identity of the synthesized compound.

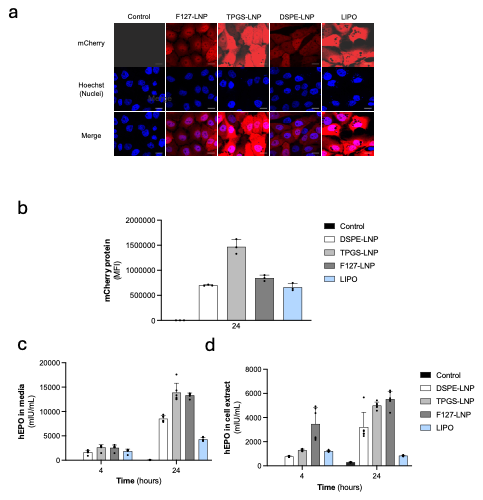

Supplemental Figure 2: (a) Representative confocal images of OK cells transfected with LNPs encapsulating mCherry mRNA. Red: presence of mCherry; blue: nucleus were stained with Hoechst 33342 (scale 15 µm). (b) In vitro mCherry mRNA expression in HKC-8 cells transfected with F127-LNP, TPGS-LNP and DSPE-LNP. (c) EPO expression in the cell media of OK cells transfected with LNPs encapsulating hEPO mRNA. (d) EPO expression in OK cells transfected with LNPs encapsulating hEPO mRNA measured by ELISA. Data are presented as Mean ± SD (n = 3); * p < 0.05, ** p < 0.01, *** p < 0.001. Significance was determined by independent t-test.

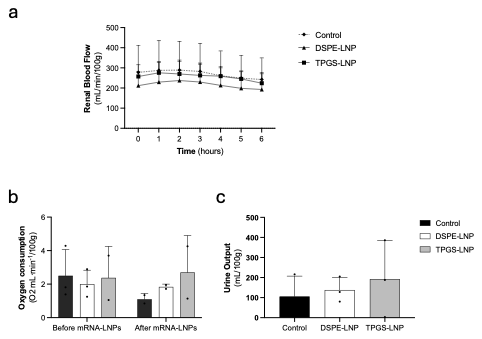

Supplemental Figure 3: Perfusion characteristics during normothermic machine perfusion (NMP) of porcine kidneys following hEPO mRNA delivery. Data are presented for the following parameters: (a) RBF (Renal Blood Flow) over time, (b) oxygen consumption (15 minutes before mRNA administration and 1 hour after mRNA administration) and (c) total urine output throughout the perfusion period. These measurements were recorded throughout the perfusion process using DSPE-LNP (carrying hEPO mRNA), TPGS-LNP (carrying hEPO mRNA) and Control (non-treated kidneys). Significance is determined by multiple Wilcoxon test for RBF, Kruskal-Wallis test for urine output and multiple two-way ANOVA for oxygen consumption. * p < 0.05, ** p < 0.01, *** p < 0.001.

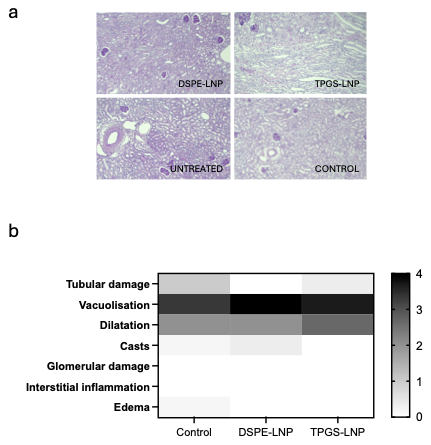

Supplemental Figure 4: Histological evaluation of kidney tissue. Formalin-fixed paraffin-embedded (FFPE) sections were stained with Hematoxylin and Eosin (H&E) and Periodic Acid–Schiff (PAS) to assess tissue morphology and structural integrity. Scoring of biopsies taken at the end of the perfusion, was performed based on predefined criteria evaluating parameters such as tubular injury, glomerular damage, and interstitial alterations. (a) Representative images and corresponding histopathological scores are shown. (b) Heatmap of quantitative histological scoring in perfused kidneys following treatment with DSPE-LNPs or TPGS-LNPs encapsulating hEPO mRNA, TPGS-LNPs without mRNA (Control) or untreated kidneys (n= 3 per group, averaged).
